# Computational design and construction of an *Escherichia coli* strain engineered to produce a non-standard amino acid

**DOI:** 10.1101/2022.04.02.486821

**Authors:** Ali R. Zomorrodi, Colin Hemez, Pol Arranz-Gibert, Terrence Wu, Farren J. Isaacs, Daniel Segrè

## Abstract

Introducing heterologous pathways into host cells constitutes a promising strategy for synthesizing nonstandard amino acids (nsAAs) to enable the production of proteins with expanded chemistries. However, this strategy has proven challenging as the expression of heterologous pathways can disrupt cellular homeostasis of the host cell. Here, we sought to optimize the heterologous production of the nsAA para-aminophenylalanine (pAF) in *Escherichia coli*. First, we incorporated a heterologous pAF biosynthesis pathway into a genome-scale model of *E. coli* metabolism, and computationally identified metabolic interventions in the host’s native metabolism to improve pAF production. Next, we explored different ways of imposing these flux interventions experimentally and found that the upregulation of flux in chorismate biosynthesis pathway through the elimination of feedback inhibition mechanisms could significantly raise pAF titers (∼20 fold) while maintaining a reasonable pAF yield-growth rate trade-off. Overall, this study provides a promising strategy for the biosynthesis of nsAAs in engineered cells.

## Introduction

Engineering microbes with diverse natural and non-naturally occurring functions is a major endeavor in synthetic biology, important for multiple goals. One goal is the microbial production of industrial and therapeutic small molecules through metabolic engineering. Another goal involves the ribosomal production of proteins containing nonstandard amino acids (nsAAs), which expand the chemistry of the canonical set of 20 amino acids that organisms in all kingdoms of life use for protein biosynthesis [1, 2]. Proteins that incorporate nsAAs enable diverse biochemistries not typically found in nature, such as hydrocarbon-based secondary structure stabilization [3], site-specific antibody-drug conjugation [4], and covalent linkage of proteins to form functional biopolymers [5, 6]. Although short nsAA-bearing proteins, such as stapled peptides, have traditionally been generated via asymmetric synthesis methods [7] and while cell-free translation techniques exist for the production of larger proteins [8], the ribosomal incorporation of nsAAs into the proteins of living cells greatly expands the scope and utility of nsAA-containing proteins. For instance, ribosomally-synthesized nsAA-containing proteins can be incorporated into biocontainment strategies that limit the survival and propagation of engineered microbes to specified environments [9, 10], fluorescent proteins that serve as *in vivo* probes for enzymatic activities [11], and sequence-defined synthetic biomaterials containing multiple instances of nsAAs [12-14].

In order to synthesize proteins that contain ribosomally-synthesized nsAAs, engineered organisms require a pool of nsAAs from which to draw during translation. To date, the problem of provisioning organisms with nsAAs has been predominantly tackled by exogenously supplementing the desired amino acids [12]. Additionally, there are only a few reports of *in vivo* production for genetic code expansion by manipulating endogenous amino acid biosynthetic pathways to favor the intracellular accumulation of intermediates that are used as nsAAs [15-17]. Both strategies are subject to limitations. In the case of exogenous supplementation, some nsAAs may have low cell membrane permeability, compromising their transport into the cell for ribosomal incorporation. In the case of native pathway production, the set of nsAAs that can be synthesized is limited to the biosynthetic capabilities of the host organism.

Incorporating and optimizing heterologous biosynthetic pathways into a host cell’s metabolism constitutes a third possible strategy for provisioning organisms with nsAAs. This circumvents the need for exogenous nsAA supplementation and enables the biosynthesis of a vast range of nsAAs beyond the host organism’s native metabolic capabilities. Heterologous pathways for synthesizing nsAAs can be obtained from bacteria, fungi, and plants that produce a wide variety of amino acids with nonstandard functional groups as intermediates when synthesizing antibiotics, toxins, and other bioactive small molecules [18]. For example, the gram-positive bacterium *Streptomyces cattleya* is capable of synthesizing a terminal-alkyne amino acid [19] and 4-fluorothreonine [20]. Importing naturally occurring biosynthetic pathways for nsAAs into model organisms with a high capacity for ribosomal nsAA incorporation, such as genomically recoded *E. coli* [1], could be an effective way to synthesize proteins using nsAAs. Despite this promise, the efficient biosynthesis of nsAAs and other small molecules via expression of heterologous pathways in laboratory host strains has proven challenging because it requires laborious efforts to mine, characterize, and optimize heterologous biosynthetic pathways in hosts, given that such pathways can disrupt cellular homeostasis by co-opting native metabolic resources for small molecules production [21, 22]. While a previous study aimed to address this challenge by engineering the native metabolism of an *E. coli* strain for producing an nsAA [23], systems-level studies of nsAA overproduction, which take into account the entire scope of cellular metabolism, are still lacking.

Genome-scale models (GEMs) of metabolism can help address this gap [24, 25]. These models consist of the full inventory of metabolic reactions encoded by the genome of an organism and can be computationally simulated to systematically explore metabolic tradeoffs in engineered organisms. A wide spectrum of computational approaches has been developed to design overproducing microbial strains by using GEMs as a basis (see [24, 25] for a review of these approaches). These tools computationally identify candidate flux changes in the network, such as knockouts, up-regulations or down-regulations that lead to the enhanced production of a biochemical of interest. The identified flux changes are then mapped to genetic manipulations that can be implemented experimentally. Several studies have reported on the successful utilization of these computational pipelines to guide the design of engineered microbial strains overproducing commodity chemicals, biofuel precursors, and a variety of other chemicals [26-30].

We hypothesized that the efficient synthesis of nsAAs using heterologous pathways also requires systematic engineering of the host’s native metabolites to proportionately allocate metabolic resources to cellular growth and the bioengineering objective, i.e., nsAA production. In this study, we tackle this by combining computational systems biology approaches based on GEMs of metabolism and synthetic biology techniques to explore the feasibility of engineering highly efficient nsAA producer microbes. We focus our efforts on optimizing the biosynthesis of *para*-aminophenylalanine (pAF) in *E. coli*, generated from intracellular chorismate using a heterologously expressed gene cluster from *Pseudomonas fluorescens [23]*. Motivated by the observation that a trade-off exists between pAF production and growth rate in this engineered *E. coli* cultured in carbon-limited environments, we sought to explore opportunities for modulating this trade-off by using computational modeling. To this end, we used a GEM of *E. coli* metabolism and a computational strain design pipeline to better understand how the introduction of the heterologous pAF-producing pathway co-opts native metabolic resources, and to computationally identify rational ways of rewiring the host metabolism to improve pAF production. We then used the predicted metabolic flux interventions as a starting point to apply multiplex genome engineering technologies [31-33] to experimentally construct and test engineered strains for pAF production. We found that upregulation of metabolic flux in the chorismate biosynthesis pathway through the elimination of feedback inhibition mechanisms is the most promising strategy to increase pAF production. However, the optimized strains continued to exhibit a trade-off between growth rate and pAF production. Our study provides a basis for the systematic exploration of host cell metabolism to optimize the biosynthesis of natural products via heterologous expression. The strategy presented here may be applied to diverse biosynthetic pathways in a wide range of host organisms.

## Results

### A trade-off between nsAA production and growth in carbon-limited environments

To enable pAF production by *E. coli*, we engineered the *E. coli* strain EcNR2 [34] to synthesize pAF using the *papBAC* gene cluster derived from *Pseudomonas fluorescens* strain SBW25. The enzymes PapA, PapB, and PapC convert chorismate into *para*-aminophenylpyruvate, which is then converted to pAF by native cellular aminotransferases [23] (**Figure 1A**). We constructed a pAF production circuit by placing the *papBAC* genes downstream of the PL_tetO_ anhydrotetracycline (aTc) inducible promoter (**Figure 1B**) and cloned the gene cluster into a plasmid with a p15A origin of replication [35]. The aTc-inducible promoter enables titratable expression of the genes under its control, a characteristic we confirmed by examining an analogous circuit with green fluorescent protein (GFP) in the place of *papBAC* [36] (**Supplementary Figure S1**).

**Figure 1.**
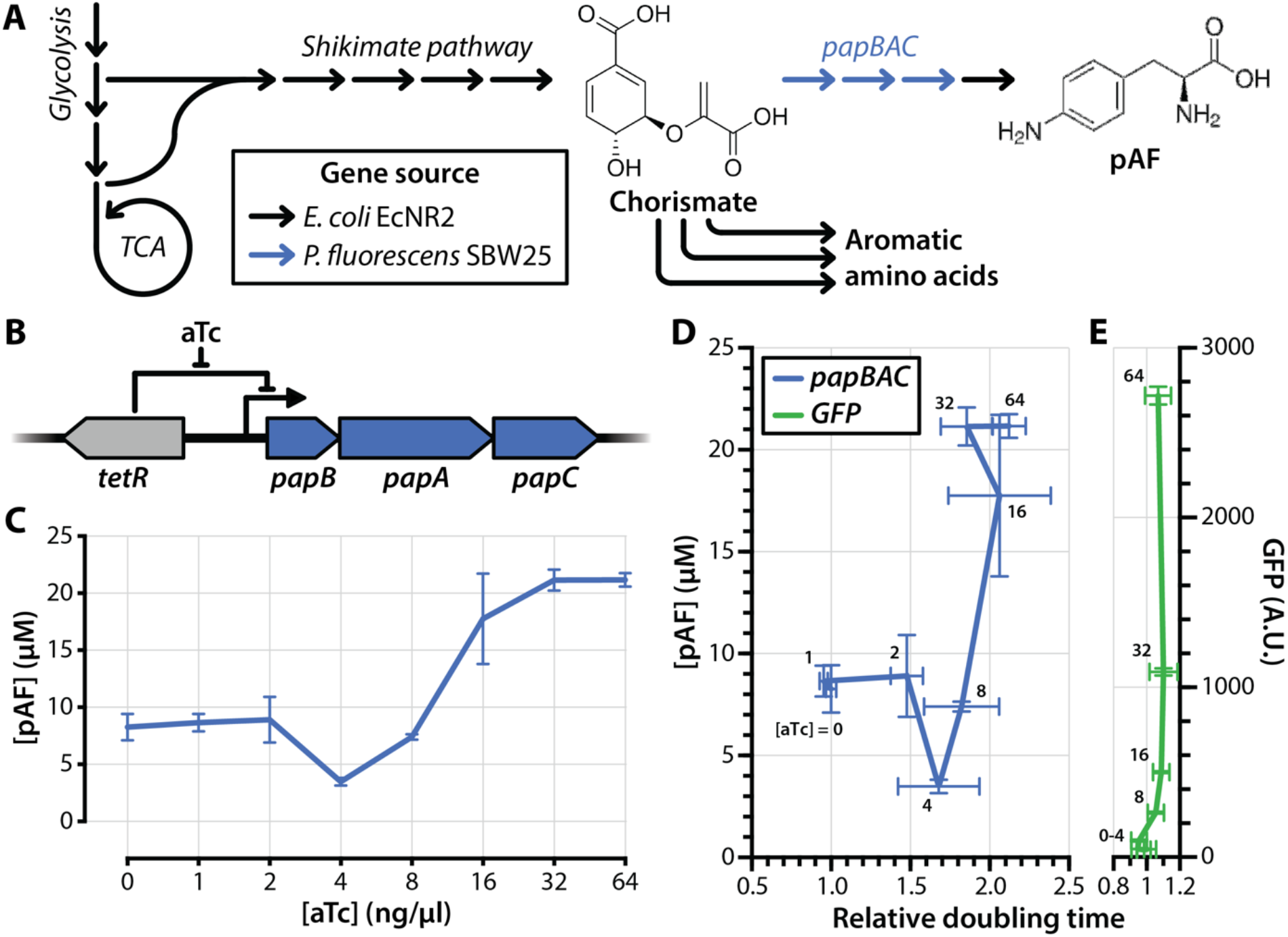
Trade-off between pAF production and growth in non-engineered strains. (**A**) Overview of pAF production via heterologous expression of *papBAC*. Chorismate, the end-product of the shikimate pathway, is converted to p-aminophenylpyruvate by the PapBAC enzymes, which is then converted to pAF by host cell deaminases. (**B**) Diagram of aTc-inducible papBAC overexpression circuit used in this study. (**C**) pAF production as a function of aTc concentration in *E. coli* strain EcNR2 bearing the aTc-inducible papBAC overexpression cassette and grown in M9 minimal medium. (**D**) pAF production and doubling time (relative to growth without aTc) for EcNR2 bearing the aTc-inducible *papBAC* overexpression cassette. (**E**) GFP fluorescence and doubling time (relative to growth without aTc) for EcNR2 bearing GFP in the place of *papBAC* under an aTc-inducible expression cassette.

We induced *papBAC* expression at a range of aTc concentrations in M9 minimal medium supplemented with 0.4% glucose and measured extracellular pAF concentrations after 24 hours of incubation using tandem mass spectrometry. We observed low levels of pAF production even in the absence of aTc, a consequence of both leaky expression under PL_tetO_ and of peak overlap in MS spectra. Concentrations of aTc above 8 ng/ml yielded pAF titers above baseline, with pAF yield plateauing at ∼20 µM (0.001 g pAF per g of glucose) beyond aTc concentrations of 32 ng/ml (Figure 1C). However, strains grown in aTc concentrations of 2 ng/ml and higher exhibited substantial increases in doubling time relative to the uninduced control (Figure 1D). Notably, the growth impairment of the strain increases concomitantly with higher *papBAC* expression, which suggests a trade-off between pAF production and growth rate. We did not observe a similar trade-off between protein expression and doubling time when we expressed non-enzymatic GFP under an analogous circuit, indicating that the trade-off observed is not due to increased translational load. We hypothesized that the trade-off is a consequence of rerouting chorismate flux toward pAF production and away from native downstream pathways, which presumably attenuate the production of metabolites essential for growth in minimal medium (Figure 1A).

### Construction of *in silico* pAF-producing *E. coli*

Our observation that pAF production trades off with growth in *E. coli* illustrates the disruption of cellular homeostasis caused by the expression of heterologous enzymatic pathways, a phenomenon observed in other related studies [21, 37-39]. This motivates the need to engineer the host cell’s native metabolism in order to optimize nsAA production in dynamic environments. To address this, we leveraged computational systems biology methods to systematically study nsAA production in *E. coli*. To construct the corresponding pAF-producing *E. coli in silico*, metabolic reactions encoded by the *papBAC* gene cluster were extracted and consolidated from several online databases including MetaCyc [40], KEGG [41] and Model SEED [42] and were incorporated into the *i*JO1266 GEM of metabolism of *E. coli* [43] (**Figure 2**). We next used this engineered metabolic model to examine the production capacity of pAF by the engineered *E. coli* strain *in silico*. This analysis revealed that this strain is able to produce a maximum of 0.54 mmoles of pAF per mmole of glucose (0.54 g pAF per g glucose) under the aerobic minimal M9 medium (equivalent of 2.16 g/L or 120 mM of pAF in an M9 medium with 4% glucose). However, this maximum is achieved at the expense of zero growth, implying that pAF production is in direct competition with growth, which was further confirmed by plotting the predicted maximum pAF production level by the model as a function of maximum growth rate (**Figure 2A**). This is due to the fact that glutamine and chorismate, which are the main precursors for the pAF biosynthesis in the network, also serve as a precursor for biomass production (*i*.*e*., growth), or for the biosynthesis of a number of other essential amino acids such as phenylalanine, tyrosine, and tryptophan (**Figure 2B**). These results suggest that further engineering of native metabolism of the host *E. coli* strain is required to enhance pAF production.

**Figure 2.**
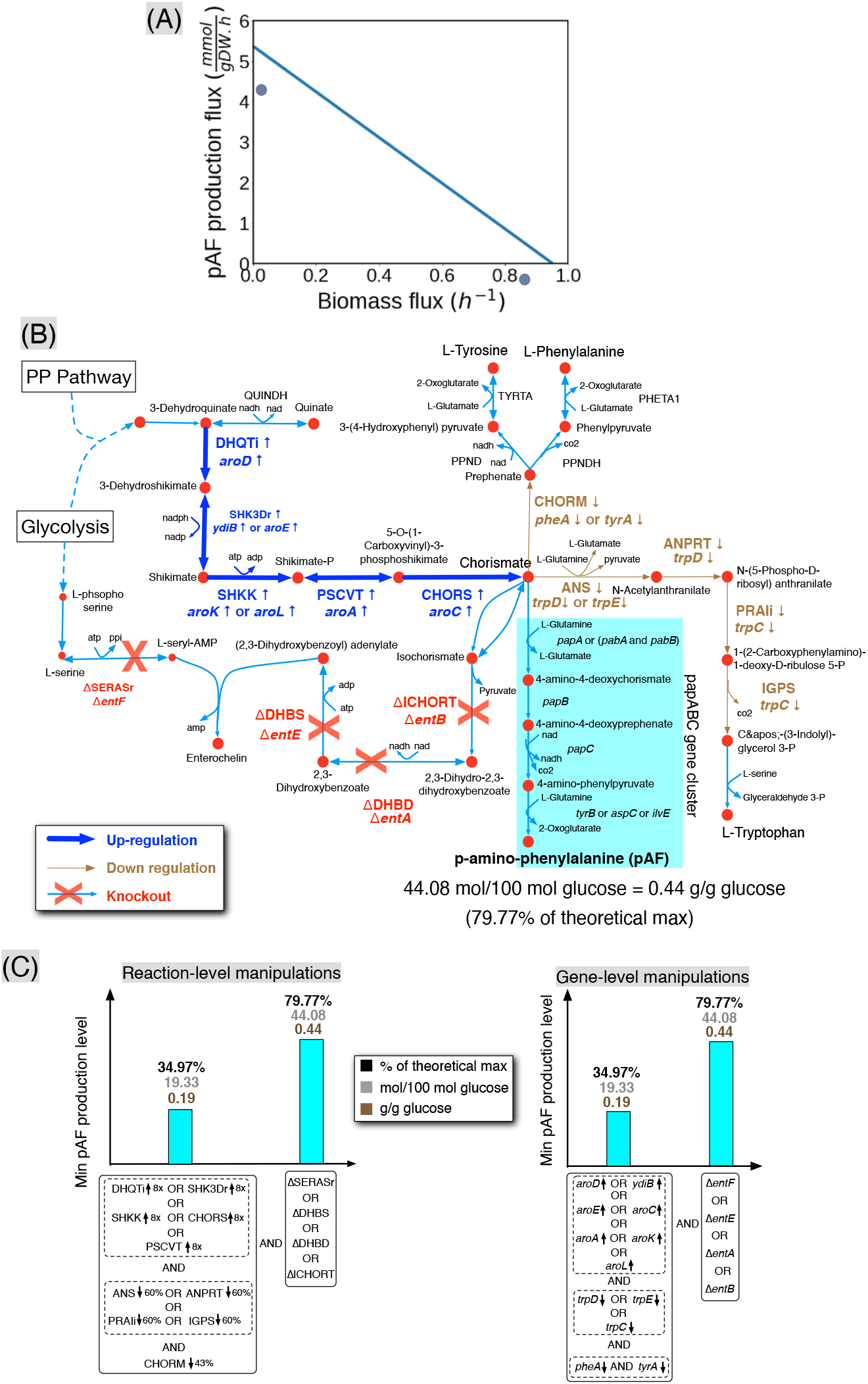
Computational design of a pAF-producing *E. coli* strain using a heterologous pathway. (A) The predicted pAF production levels as a function of the biomass production flux (*i*.*e*., growth). (B) Identified metabolic interventions to enhance pAF production. (C) The impact of predicted metabolic flux interventions (left panel) on the minimum (*i*.*e*., guaranteed) pAF production in the metabolic network and the corresponding gene-level interventions inferred from gene-reaction associations in the model (right panel). A minimum of three reaction (or gene) interventions are needed to achieve a non-zero minimum pAF production yield.

### Computational design of metabolic interventions for the native metabolism of the host *E. coli* strain

We used the computational strain design pipeline OptForce [44] (Methods) to identify metabolic interventions leading to the improved production of pAF in the *E. coli* strain harboring the *papBAC* gene cluster under the aerobic minimal M9 medium using glucose as the carbon source. The first set of interventions that we identified consists of the upregulation of reactions encoded by the *papBAC* gene cluster that results in a minimum pAF production of ∼80% of theoretical maximum when the growth is set to be at least 20% of maximum. These reaction flux increases draw the metabolic flow toward pAF and can be also readily inferred intuitively. However, pull strategies alone may not always be sufficient to provide a desired production level for a target biochemical due to the presence of bottlenecks in other parts of the metabolic network, which cannot be directly captured by GEMs of metabolism. Therefore, in the next step, we sought to identify metabolic interventions in other parts of the host’s native metabolism that push metabolic flow toward pAF. This was done by re-running OptForce while preventing the upregulation of reactions encoded by the *papBAC* gene cluster. This analysis identified a set of reaction manipulations that collectively lead to an increased pool of chorismate (**Figure 2B**). Here, we found that at least three simultaneous reaction manipulations are required to enable a minimum pAF production based on the model. These reaction manipulations result in a minimum pAF production level of 34.97% of theoretical maximum (**Figure 2C**) and include (i) the increasing flux in any of the reactions in the shikimate pathway that directly lead to the biosynthesis of chorismate (encoded by *aro* family genes or *ydiB*) (Figure 2B); (ii) decreasing flux in any of the reactions that convert chorismite to aromatic amino acids namely L-tryptophan (encoded by *trpC, D* and *E*), and to phenylalanine and tyrosine (chorimsate mutase “CHORM”, encoded by *pheA* or *tyrA*) (Figure 2B). In addition, we found that a fourth intervention, *i*.*e*., the deletion of any of the reactions encoded by *entA, entB, entE* and *entF* to prevent the conversion of chorismate to enterochelin, will increase the minimum pAF production more than twofold (from 34.97% to 79.77% of theoretical maximum) (**Figure 2C**). Although the deletion of aromatic amino acid genes to improve pAF production has been explored before [23], such modifications would lead to auxotrophies and other interventions identified in this study have not been reported, thus highlighting the importance of taking into account the entire scope of metabolism (afforded GEMs) to infer potentially promising metabolic interventions while minimizing impact on growth and fitness.

It is worth noting that the analysis of epistatic interactions among our identified flux manipulations with respect to pAF production as the phenotype of interest revealed that (**Supplementary Figure S1**) no single or pair combinations of the four reaction manipulations noted above (**Figure 2B**) is enough to enable a non-zero minimum pAF production in the network. Of note, an epistatic interaction between two genes means that the phenotype of a double-gene mutant strain cannot be easily inferred from phenotypes of single-gene mutant strains [45]). However, with simultaneous implementation of these four reaction manipulations, a high metabolic flux is pushed toward chorismate while all metabolic sinks of chorismate (except those that are essential for the synthesis of growth precursors) are blocked, thereby funneling a high carbon flux toward pAF. This high nonlinearity highlights the importance of computational analyses to identify interventions that may need to be applied concurrently in order to have a significant effect.

### Construction and characterization of computationally designed overproducing strains

Computational modeling identified numerous opportunities for metabolic flux manipulations to enhance pAF production in *E. coli*, including the downregulation of metabolic flux downstream of chorismate biosynthesis and the upregulation of flux upstream of chorismate biosynthesis. These predictions serve as a starting point to experimentally design an engineered *E. coli* strain with an enhanced pAF production capacity. Although we can implicitly infer genetic manipulations corresponding to predicted flux interventions using gene-reaction associations in GEMs (**Figure 2C**), there can be different ways of imposing these flux changes experimentally, not all of which could be directly captured by GEMs. For example, increasing metabolic flux in the shikimate pathway (toward chorismite) can be achieved experimentally by either upregulating the *aro* family genes or by removing feedback inhibitions in this pathway (even though the latter is not directly captured by stoichiometric models used in this study) (**Figure 2B**). In particular, the redundant 3-deoxy-7-phosphoheptlonate synthases AroF, AroG, and AroH are allosterically inhibited by tyrosine, phenylalanine, and tryptophan, respectively [46-48], and the transcriptional repressor TyrR reduces the expression of numerous *aro* family genes under conditions of tyrosine abundance [49]. Thus, we hypothesized that these feedback mechanisms could play a strong role in enhancing or attenuating metabolic flux toward chorismate synthesis. Therefore, we sought to eliminate these inhibition loops in our engineered strains as a way of increasing metabolic flux toward chorismate predicted by computational studies.

We implemented model-guided metabolic flux downregulations by knocking out genes downstream of chorismate biosynthesis (*pheA, trpDE, tyrA*, and *entA*), and flux upregulations by cloning an additional copy of the penultimate gene upstream of chorismate biosynthesis (*aroC*) into episomal expression vectors. We used multiplex genome engineering and recombineering [50] to construct *E. coli* strains with combinatorial knockouts of *pheA, trpDE, tyrA*, and *entA*, genes downstream of chorismate biosynthesis. The knockout of aromatic amino acid biosynthesis genes generates auxotrophies for the respective amino acids. To compensate for these auxotrophies, we grew engineered strains in minimal medium supplemented with phenylalanine, tryptophan, and tyrosine. We also cloned *aroC* and placed it under the control of a vanillic acid-inducible promoter (*P*_*vanR*_) [51] to enable titratable overexpression. To verify whether genes under the control of the two synthetic promoters PL_tetO_ and *P*_*vanR*_ could be induced independently of one another, we also constructed a circuit analogous to the one used for *papBAC* and *aroC* expression with GFP under the control of P_tetO_ and RFP under the control of *P*_*vanR*_ (**Supplementary Figure S2**). We observed that GFP expression was unaffected by vanillic acid concentrations of up to 4 µM and that RFP expression was unaffected by aTc concentrations of up to 32 ng/ml (**Supplementary Figure S3**). Finally, we used MAGE [31] to introduce nonsynonymous point mutations that render AroF and AroG allosterically insensitive to tyrosine and phenylalanine, respectively [46, 47]. Because AroF and AroG together account for >99% of 3-deoxy-7-phosphoheptlonate synthase activity within metabolically active *E. coli* [52], we did not introduce analogous feedback-inactivating mutations to AroH.

We investigated the effects of these interventions, alone and in combination, on pAF production after 24 hours of induction in minimal media supplemented with tyrosine, phenylalanine, and tryptophan (**Figure 3A**). Titration of vanillic acid to overexpress *aroC* led to modest increases in pAF production, while no combination of downstream gene knockouts increased pAF yields contrary to our expectation. Feedback-inhibitory mutations did not lead to increases in pAF production on their own, but we observed a > 4-fold increase in pAF production in strains with both AroFG feedback inhibition mutations (*aroFG*-FIM) and a *tyrR* knockout (Δ*tyrR*). This suggests the existence of redundant feedback control mechanisms within the shikimate pathway that regulate flux toward chorismate, leading to the observed epistatic pattern on pAF production.

**Figure 3.**
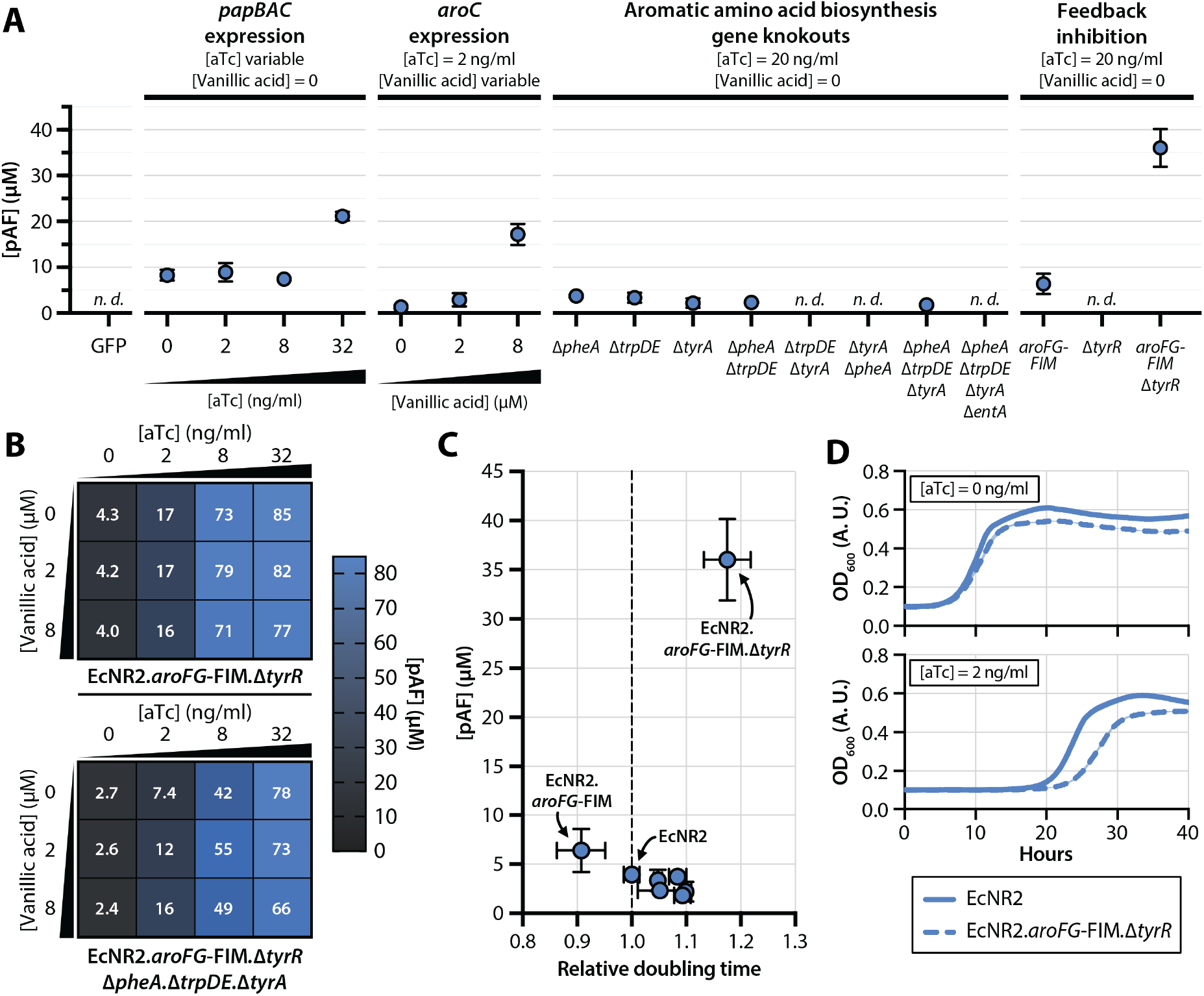
pAF production and growth rate trade-offs in engineered strains. (**A**) pAF production in strains with modulations in aTc-induced *papBAC* overexpression, vanillic acid-induced *aroC* overexpression, aromatic amino acid biosynthesis gene knockouts, and shikimate pathway feedback inhibitions. “GFP” condition denotes a strain bearing an aTc-inducible GFP overexpression cassette. All strains were grown in M9 minimal medium supplemented with phenylalanine, tryptophan, and tyrosine. (**B**) pAF production in strains with shikimate pathway feedback inhibition mutations (*aroFG*-FIM and Δ*tyrR*) and modulated *papBAC* and *aroC* overexpression. The strain in the top panel bears feedback inhibition mutations only, and the strain in the bottom panel bears knockouts in *pheA, trpDE*, and *tyrA* in addition to feedback inhibition mutations. (**C**) pAF production and doubling time (relative to the growth of EcNR2) for strains with different combinations of genomic interventions. Strains in which pAF production was not detected are not included. (**D**) Representative growth curves of EcNR2 and EcNR2.*aroFG*-FIM.Δ*tyrR* bearing aTc-inducible *papBAC* overexpression cassettes at 0 aTc (top panel) and 2 ng/µl aTc (bottom panel).

We hypothesized that other epistatic patterns may exist among the genomic interventions targeted, and we reassessed the effects of gene overexpression and knockout in the context of a feedback-insensitive strain. Tuning the expression of the *papBAC* genes through aTc titration had a strong effect on pAF yield in ECNR2.*aroFG*-FIM.Δ*tyrR*, up to a maximal pAF yield of 85 µM after 40 hours of induction (0.004 g pAF per g of glucose) (**Figure 3B**, top panel). *aroC* overexpression via titration of vanillic acid led to no increases in pAF yield when paired with any level of *papBAC* induction. The expression of *aroC* was in fact detrimental to pAF yield at high aTc concentrations, possibly because of the high translational load placed on cells expressing both AroC and PapB/A/C in abundance. These same patterns—of pAF production driven predominantly by *papBAC* gene overexpression and of detrimental effects of *aroC* expression on pAF yield—were also observed in feedback-insensitive strains with additional knockouts in *tyrA, pheA*, and *trpDE* (**Figure 3B**, bottom panel).

### Relationship between pAF production and growth in engineered strains

Given our initial observation of a trade-off between pAF production and growth rate (**Figure 1D**), we sought to determine how the model-guided genomic interventions (**Figure 3A**) influence the balance between cellular growth and pAF production in *E. coli*. Our efforts to culture engineered strains with *tyrA, pheA*, and *trpDE* knockouts in 96-well plate readers under conditions of *papBAC* induction were unsuccessful, presumably because suboptimal agitation and aeration conditions in such instruments inhibits the growth of strains with fitness-reducing gene knockouts. We thus measured the growth rates of all strains in the absence of *papBAC* expression (aTc = 0) and compared these rates against pAF yields obtained from the same strains in conditions optimal for pAF production (see Methods) (**Figure 3C, Supplementary Table S1**). Most engineered strains exhibited longer doubling times than EcNR2 without improving pAF yields. EcNR2.*aroFG*-FIM showed a modestly lower doubling time (0.91 ± 0.04 relative to EcNR2) and modestly higher pAF yields (6.4 ± 2.2 µM), while EcNR2.Δ*tyrR* showed a longer doubling time (1.14 ± 0.03 relative to EcNR2) and no detectable pAF yield.

EcNR2.*aroFG*-FIM.Δ*tyrR* was nonetheless capable of growing in a plate reader under *papBAC* induction conditions. Compared to EcNR2, the strain showed a 18% slower growth rate (161 ± 5 min for EcNR2.*aroFG*-FIM.Δ*tyrR* versus 137 ± 1 min for EcNR2) in the absence of *papBAC* expression (**Figure 3D**, top panel). However, both EcNR2 and EcNR2.*aroFG*-FIM.Δ*tyrR* exhibited profound lag phases at low levels of *papBAC* expression ([aTc] = 2 ng/ml; **Figure 3D**, bottom panel). This suggests that feedback inhibition mutations can increase pAF yields but do not resolve the trade-off between pAF production and growth rate in carbon-limited media.

## Discussion

The optimization of heterologous pathways for nsAA production is a promising strategy for engineering microbes with diverse biochemical capabilities. Heterologous expression often places significant metabolic burdens on engineered strains, which may hinder their utility in diverse industrial, biomedical, and environmental contexts. In this study, we leveraged GEMs of metabolism to guide the design of *E. coli* strains capable of producing the nsAA pAF from a heterologous pathway with minimal disruptions to cellular homeostasis. Previous studies have successfully engineered native *E. coli* metabolism to maximize pAF production in fermentation settings [23], wherein the active growth of cells is of minimal concern, and did not take into account the entire metabolism of the host. We build on this prior work by simultaneously accounting for growth and pAF production in our strain engineering efforts and by considering the entire scope of the host’s native metabolism using GEMs. This enables us to systematically pinpoint and test the effects of metabolic interventions that span the entire *E. coli* metabolome.

Our initial efforts for pAF production in non-engineered *E. coli* hosts were hindered by a conspicuous trade-off between pAF production titers and growth rate (**Figure 1D**), presumably because expression of the pAF-producing *papBAC* genes disrupts cellular homeostasis by shunting essential metabolites away from central metabolism. Computational modeling using GEMs identified numerous metabolic flux interventions that were predicted to improve the production of pAF for a given minimal growth rate. These interventions included decreasing metabolic flow downstream of chorismate biosynthesis through the downregulation (or knockout) of genes *tyrA, pheA, trpDE*, and *entA* and increasing metabolic flux upstream of chorismate biosynthesis through the upregulation of genes in the *aro* family (such as *aroC)*. GEMs of metabolism, which are based on a stoichiometric representation, do not account for the nonlinear effects of regulatory interactions. Therefore, we also decided to consider the elimination of feedback inhibition mechanisms within the shikimate pathway as an alternative to implement flux upregulations to increase metabolic flow towards chorismite as predicted by using GEMs of metabolism. Although the removal of genes involved in the aromatic amino acids biosynthesis (more specifically pheA) has been implemented before to increase pAF production [23], their combination with other interventions we identified in this study has not previously been explored.

Interestingly, we found that increased metabolic flow in the shikimate pathway through *aroC* upregulation, either alone or in combination with other predicted metabolic interventions, did not lead to increased pAF production compared to the non-engineered host (**Figure 3A**). This could be due to the presence of other rate limiting enzymatic reactions upstream of AroC in the shikimate pathway, implying that additional gene upregulations in that pathway are needed to impose the desired effect. However, we observed that imposing an increased metabolic flow toward chorismite through the elimination of feedback inhibition loops (via the elimination of allosteric feedback on AroF and AroG and deletion of the tyrosine-sensitive transcriptional repressor TyrR) led to a measurable increase in pAF production. Contrary to our expectation, combining these feedback elimination interventions with other computationally predicted interventions did not improve pAF production any further. These observations suggest that specific genetic interventions inferred directly from gene-reaction associations in the GEMs of metabolism may not always behave as expected due to regulatory/allosteric effects that are not captured by these models. This alludes to the need to consider additional regulatory interactions in these pathways in future engineering efforts, *e*.*g*., by using large-scale kinetic models of metabolism [53, 54] (instead of stoichiometric models used in this study), which can take into account such regulatory and allosteric interactions.

The interventions that did increase pAF production in our engineered strain, AroFG-FIM and Δ*tyrR*, showed an epistatic effect on pAF production. Neither AroFG feedback elimination nor *tyrR* knockout raised pAF titers when implemented in isolation but installing both interventions within the same strain led to substantial increases in pAF titer (**Figure 3A**). This epistatic pattern could be due to redundant regulatory mechanisms within the shikimate pathway that control chorismate production. Both AroF and TyrR respond to high tyrosine conditions by reducing flux through the shikimate pathway, and the elimination of feedback nodes on both genes is thus needed to restore chorismate production in the tyrosine-supplemented minimal media used in this study. We observed substantial (∼20-fold) increases in pAF production in strains with engineered chorismate pathway regulation compared to unmodified strains (**Figure 3B**). In future studies, we propose that it is possible to achieve further increases in pAF production through more extensive metabolic feedback loop modulation, but that these interventions do not necessarily alter the trade-off between pAF production and growth (**Figure 3D**).

Sourcing nonstandard amino acids through biosynthesis *in vivo* holds promise for more efficient and cost-effective production of synthetic proteins and biomaterials. It could also serve as mediators of cellular signaling or the establishment of synthetic cross-feeding channels between members of engineered microbial communities. More specifically, we envision the exchange of nsAAs between organisms capable of producing them and organisms dependent upon them for survival as a promising approach to constructing stable microbial consortia that are minimally affected by surrounding inter-species metabolite exchanges. Engineered cross-feeding channels may also be resilient to invasion by natural strains through the exchange of metabolites that are not a component of natural cellular milieu. Crucially, organisms participating in such cross-feeding channels would need to both produce nsAAs and grow robustly in dynamic environments. Engineering microbial consortia thus requires the simultaneous optimization of two phenotypes—growth and metabolite production.

## Limitations of this study

Our study identified three classes of genomic interventions that could influence pAF production in *E. coli*: (1) the knockout of genes downstream of chorismate biosynthesis, (2) the upregulation of genes upstream of chorismate biosynthesis, and (3) the elimination of regulatory and allosteric feedback mechanisms that reduce chorismate biosynthesis in conditions of high aromatic amino acid abundance. While in this study we aimed to explore all these interventions both individually and in combination, we only explored one intervention of the second class—the overexpression of *aroC*, which encodes the final enzyme in chorismate biosynthesis. We did not observe higher pAF titers with *aroC* overexpression, which highlights the possibly that there may exist other rate-limiting reactions upstream of *aroC* that limit flux through the shikimate pathway not explored in our studies. The systematic overexpression of genes upstream of chorismate biosynthesis could reveal these rate-limiting reactions and lead to increased pAF production.

As noted earlier, stoichiometric GEMs of metabolism do not capture interventions of the third class—regulatory and allosteric feedback mechanisms. This limits their capability for predicting promising metabolic intervention strategies. We did observe increases pAF production with the inactivation of these feedback mechanisms, namely by eliminating allosteric regulation of AroF/G/H and knockout of the transcriptional repressor *tyrR* (**Figure 3A**). However, feedback mechanisms are difficult to model computationally and to tune experimentally—for instance, tuning the allosteric response of AroF/G/H would require protein sequence alterations, whose effects are more difficult to predict than genetic up- or downregulations. Although our study highlights feedback modulation as a promising strategy for the biosynthesis of nsAAs in heterologous hosts, the generalizability of this approach is limited in the absence of synthetic biology tools to control these feedback mechanisms.

## Conclusions

In this study, we integrated computational systems biology modeling using GEMs of metabolism and synthetic biology techniques to construct an *E. coli* strain with enhanced pAF production. We found that the elimination of feedback inhibition loops in the chorismate biosynthesis pathway has the highest impact on enhancing pAF production while minimizing the disruption of cellular homeostasis and growth. Imposing other interventions in pathways downstream of chorismate, which was predicted by our computational models and seemed promising, did not lead to a further increase in pAF production contrary to our expectation. However, these observations warrant future studies that consider tunable regulatory circuits in this part of the metabolism to design a more efficient pAF-producing strain. Overall, our study sets a basis for the engineering of overproducing strains, which can be used in synthetic microbial consortia that meet the dual objectives of high-level product formation and sustained growth. We anticipate that this approach could be applied to heterologous pathways beyond pAF production such as other nsAAs or biochemicals.

## Methods

### Computational methods

#### OptForce algorithm

In brief, OptForce [44], first characterizes the phenotypic space of a reference (wild-type) strain using any available metabolic flux data by minimizing and maximizing the flux of each reaction in the network subject to these experimental flux data. It next identifies the phenotypic space of an overproducing strain in a similar fashion by imposing a constraint on the minimum required biomass production (growth) in the network as well as a desired production level for the target product. By super-imposing the flux ranges in the wild-type and overproducing strains for each reaction, the list of reactions whose fluxes must increase, decrease, or shrink to zero in order to achieve the target production level of the desired product is identified. This list of reactions is then used to identify a minimal set of direct metabolic flux interventions (upregulations, downregulations, or knockouts) to achieve a specific production level for the biochemical of interest. In doing so, OptForce simulates a worst-case scenario where the network fights against imposed interventions by maximizing the minimum product formation. OptForce has been previously used successfully to design overproducing strains [28, 29].

#### Details of OptForce implementation

To characterize the phenotypic space of the wild-type strain, we used experimental flux data for 35 reactions in the central metabolism of *E. coli* from a previous study [29]. We performed the OptForce simulations by requiring a biomass production of at least 20% of theoretical maximum, for which the maximum achievable pAF production is predicted to be 44.27 (∼80% of theoretical maximum). Therefore, we set 80% of theoretical maximum as our desired pAF production level. In our analyses, reactions with no gene association in the metabolic model were prevented from any type of manipulation. Furthermore, reactions associated with *in silico* or *in vivo* essential genes in the minimal M9 medium were prevented from being removed. Re-running the OptForce with different thresholds for growth and pAF production did not affect the set of identified metabolic interventions. All simulations were performed in Python using customized scripts (Supplementary Software S1). The OptForce optimization problem was solved using the Gurobi solver (https://www.gurobi.com) in Pyomo, an optimization modeling environment in Python [55].

### Experimental methods

#### Materials, strains, and media

All strains in this study are derived from *Escherichia coli* EcNR2 [31]. LB min media from AmericanBio (Canton, MA) was used for routine strain growth and cloning. Growth and pAF production assays were performed in M9 minimal medium supplemented with 0.25 µg/L betaine and vitamin and trace mineral mixes from Neidhardt EZ rich defined medium [56] (see **Supplementary Table S2** for media components). Where indicated, M9 medium was also supplemented with 0.2 mM tyrosine, 0.4 mM phenylalanine, and 0.1 mM tryptophan to support the growth of strains with knockouts in aromatic amino acid biosynthesis genes. 0.4% (w/v) glucose was used as a carbon source for all M9 media formulations. 50 µg/ml carbenicillin was used for the growth of EcNR2 and derivatives in all conditions. Strains harboring *papBAC, aroC*, and GFP expression plasmids were cultured in carbenicillin and 95 µg/ml spectinomycin.

#### Cloning and strain engineering

Synthetic DNA fragments encoding codon-optimized *papA, papB*, and *papC* genes from *Pseudomonas fluorescens* were synthesized by GenScript (Piscataway, NJ). Primers and single-stranded oligonucleotides used for cloning and MAGE were synthesized by Integrated DNA Technologies (Coralville, IA) (**Supplementary Table S3**). Synthetic inducible circuits were generated by cloning DNA fragments into linearized plasmids with a p15A origin of replication and a spectinomycin resistance cassette. Linear DNA fragments were amplified using high-fidelity PCR kits from Kapa Biosystems sourced from Millipore Sigma (Burlington, MA). Gibson assembly (New England Biolabs; Boston, MA) was used for plasmid assembly.

MAGE [31] and recombineering [50] were used to implement all genomic perturbations described in the study. Small genomic interventions (i.e., *aroF* and *aroG* feedback inhibition mutations) were implemented by performing 3 rounds of MAGE using 90-mer single-stranded DNA oligonucleotides encoding the desired mutation. Cultures were then plated on LB agar to isolate individual clones, which were then screened for the desired mutation(s) using multiplex allele-specific colony PCR [57]. For scarless gene-scale deletions, we first recombineered the counter-selectable marker *tolC* [58] into the locus to be deleted and selected for recombinants by plating on LB + 0.1% sodium dodecyl sulfate. We then used MAGE to delete *tolC* and selected for scarless mutants by growing cultures for 8 hours in LB containing colicin E1. Cultures passing the liquid selection were then plated on LB agar to isolate individual clones. All genomic interventions were then confirmed via Sanger sequencing.

#### Growth assays

Cells were plated for single colonies on LB agar containing the appropriate antibiotics and grown overnight. Replicate (n = 3) single colonies were picked and inoculated into 3 ml LB min broth with the appropriate antibiotics. Once cell cultures reached mid-log phase (OD_600_ 0.4-0.5), they were diluted 1:100 into 3 ml M9 media without inducers. Cells were again grown to mid-log, OD-normalized to 0.4, and washed 1x via centrifugation with M9 medium to remove residual LB medium. 3 µl of OD-normalized culture was then inoculated into 147 µl M9 with appropriate inducers in a 96-well bioassay plate. Cultures were grown with shaking at 34 °C in a Biotek Synergy HT microplate reader and OD_600_ was measured every 10 minutes. Growth curves were analyzed using MATLAB scripts written to calculate doubling times (Supplementary Software S2).

#### pAF production assays

Cells were plated for single colonies on LB agar containing the appropriate antibiotics and grown overnight. Single colonies were picked and inoculated into 3 ml LB min broth with the appropriate antibiotics. Upon reaching mid-log phase, cultures were OD-normalized to 0.4 and washed 1x via centrifugation with M9 medium to remove residual LB medium. OD-normalized cultures were diluted 1:100 into 3 ml of fresh M9 medium with supplements (see “Materials, strains, and media”). Cells were grown at 34 °C with shaking for 5 hours, and *papBAC* and *aroC* were induced with the addition of aTc and vanillic acid, respectively, to the cultures. Cells were incubated at 34 °C with shaking for 24 or 40 hours. After the induction period, cultures were centrifugated and supernatant was collected for analysis via LC/MS.

### Sample Preparation and Liquid Chromatography/ Mass Spectrometer analysis (LC/MS analysis)

Supernatant from bacterial cell cultures was collected. To precipitate protein, the supernatant was treated with a 1:10 volume of 100% (w/v) ice-cold TCA, incubated on ice for 20 minutes, then centrifuged at 17,000 x g for 10 mins. This supernatant contained the compounds of interest. Samples were injected onto an Agilent Eclipse Plus C18 RRHD column (2.1×50mm, 1.8um particle size) and run over a gradient of 8 minutes. The mobile phase consisted of a linear gradient of (A) 0.1% formic acid in water and (B) 0.1% formic acid in acetonitrile using LCMS grade solvents (Optima, Fisher Chemical). The gradient parameters were as follows: 0-1 min, 10% B (isocratic wash); 1-8 min, 10-95% B;8-10 min, 95-10% B. The flow rate throughout the procedure was 0.5 ml/min. Injection volume was 12ul. The source gas temperature was set at 280 C at 11L/min, sheath gas was set to 350 C at 11 L/min. Nebulizer was set at 40 psig. Positive ion mode capillary voltage was 5.5 kV, and the nozzle voltage set at 2.0 kV. Separation and analysis were performed using an Agilent 1290 Infinity UPLC system coupled to an Agilent 6550 QToF iFunnel mass spectrometer in ESI+ mode. Data was collected in ms mode, scan range of 110 -1700, at 3 spectra/sec Lock mass spray was employed for all analysis. pAF concentration was quantified using standards consisting of known concentrations of pAF dissolved in supernatant of EcNR2 grown in M9 minimal medium (**Supplementary Figure S4**). Reference standards were generated in technical triplicate for each LC/MS experiment. A standard curve was generated from extracted ion chromatograms using the mean of the area under the curve of reference standards to interpolate pAF concentration in biological samples (**Supplementary Figure S5**).

## Supporting information

Supplementary Software S2

## Acknowledgements

We gratefully acknowledge funding from the Defense Advanced Research Projects Agency (Purchase Request No. HR0011515303, Contract No. HR0011-15-C-0091) and The Dutch Research Council (NWO). DS also acknowledges funding from the Kilachand Multicellular Design Program, the U.S. Department of Energy, Office of Science, Office of Biological & Environmental Research through the Microbial Community Analysis and Functional Evaluation in Soils Science Focus Area Program (m-CAFEs) under contract number DE-AC02-05CH11231 to Lawrence Berkeley National Laboratory, and the NSF Center for Chemical Currencies of a Microbial Planet (C-CoMP, publication #005). We also thank members of the Segrè lab and Paul Muir and Koen Vanderschuren in the Isaacs lab for their invaluable feedback on this work.

## Supplementary files

**Supplementary Figure S1:** Epistatic interactions among computationally identified flux manipulations with respect to pAF production as predicted by GEMs of metabolism.

**Supplementary Figure S2:** Circuit showing *papBAC* and *aroC* induction, titratable expression of each independently and together using GFP and RFP.

**Supplementary Figure S3:** GFP and RFP expression under the dual aTc- and vanillic acid-inducible promoter circuit.

**Supplementary Figure S4:** Representative extracted ion chromatograms for detection of pAF by LC/MS.

**Supplementary Figure S5:** Standard curve generated for pAF quantification by LC/MS.

**Supplementary Table S1:** Doubling times and pAF yields for Figure 3C.

**Supplementary Table S2:** Composition of the supplemented M9 growth medium used in experiments.

**Supplementary Software S1:** Python scripts for implementing OptForce are available at https://github.com/zomorrodilab/publications_SI/tree/main/pAF_nsAA.

**Supplementary Software S2:** MATLAB script for calculating doubling times from 96-well microplate growth curve data.

