## Supplementary Figures S1 to S5 for "Computational design and construction of an *Escherichia coli* strain engineered to produce a non-standard amino acid"

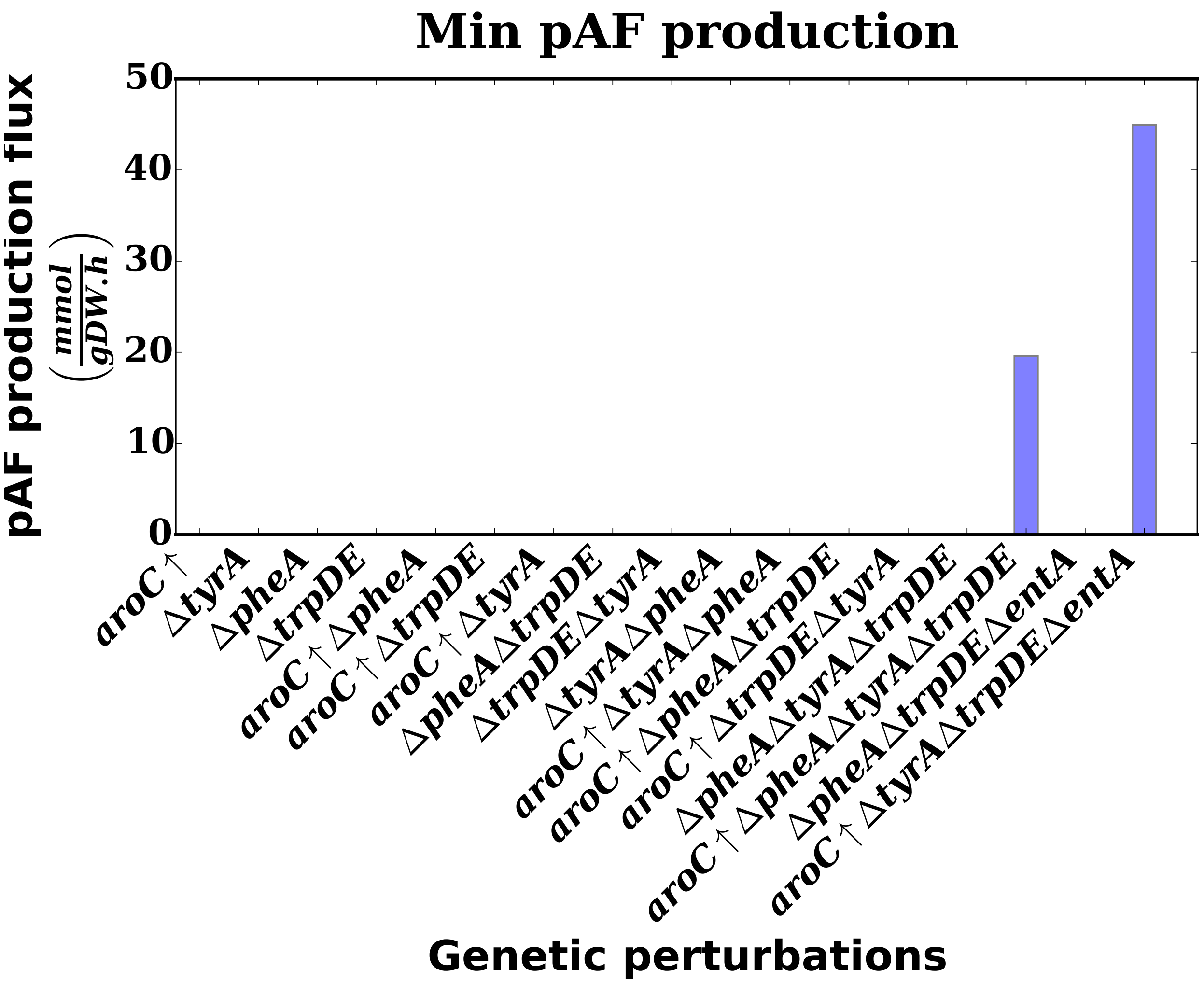


**Figure S1**. Epistatic interactions among computationally identified flux manipulations with respect to pAF production as predicted by GEMs of metabolism.

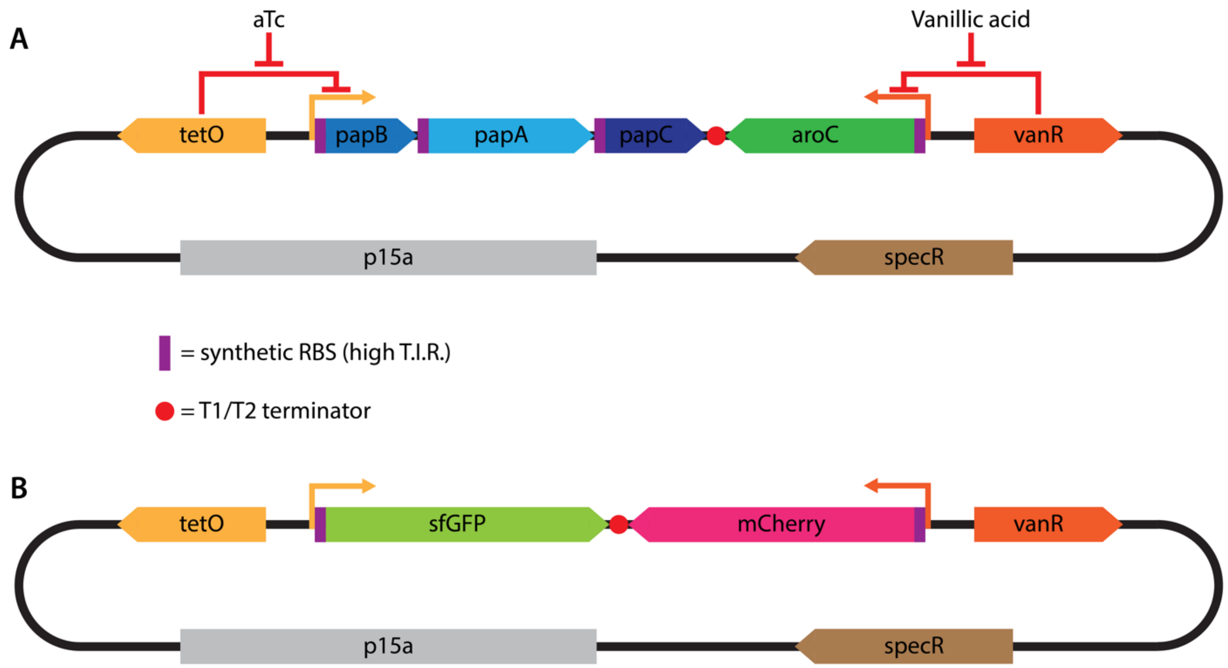


**Figure S2**. Overexpression of *papBAC* and *aroC* using dual titratable promoters. (A) Schematic of *papBAC/aroC* overexpression circuit (pPAF), with *papBAC* expressed using the anhydrotetracycline-inducible *pLtetO* promoter and aroC expressed using the vanillic acid-inducible pVanR promoter. (B) Schematic of control circuit (pPF) expressing sfGFP in the place of papBAC and mCherry in the place of aroC. T.I.R.: Translation initiation rate.

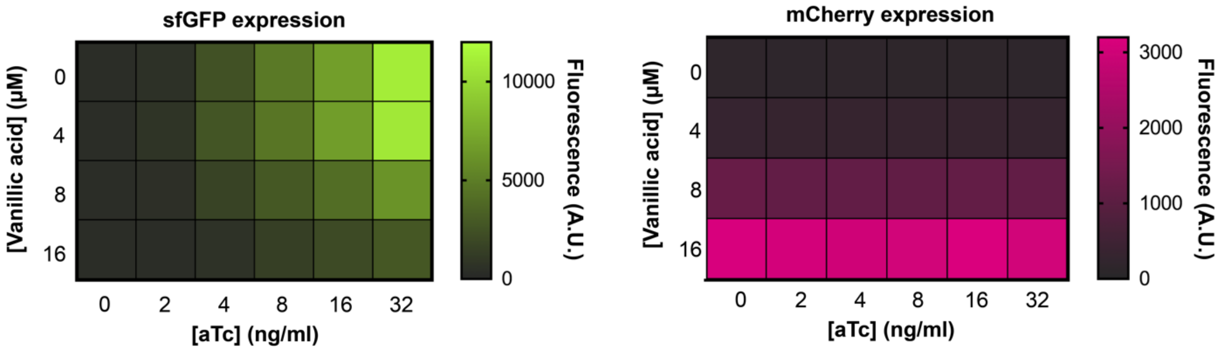


**Figure S3**. Gene expression with dual titratable promoters. EcNR2 harboring plasmid pFP (Supplementary figure S1B) was titrated with aTc to induce sfGFP expression and vanillic acid to induce mCherry expression. OD-normalized fluorescence at each combination of inducer concentrations was measured in triplicate at 10-minute intervals on a 96-well microplate reader; the maximum fluorescence measured for both sfGFP and mCherry is shown above.

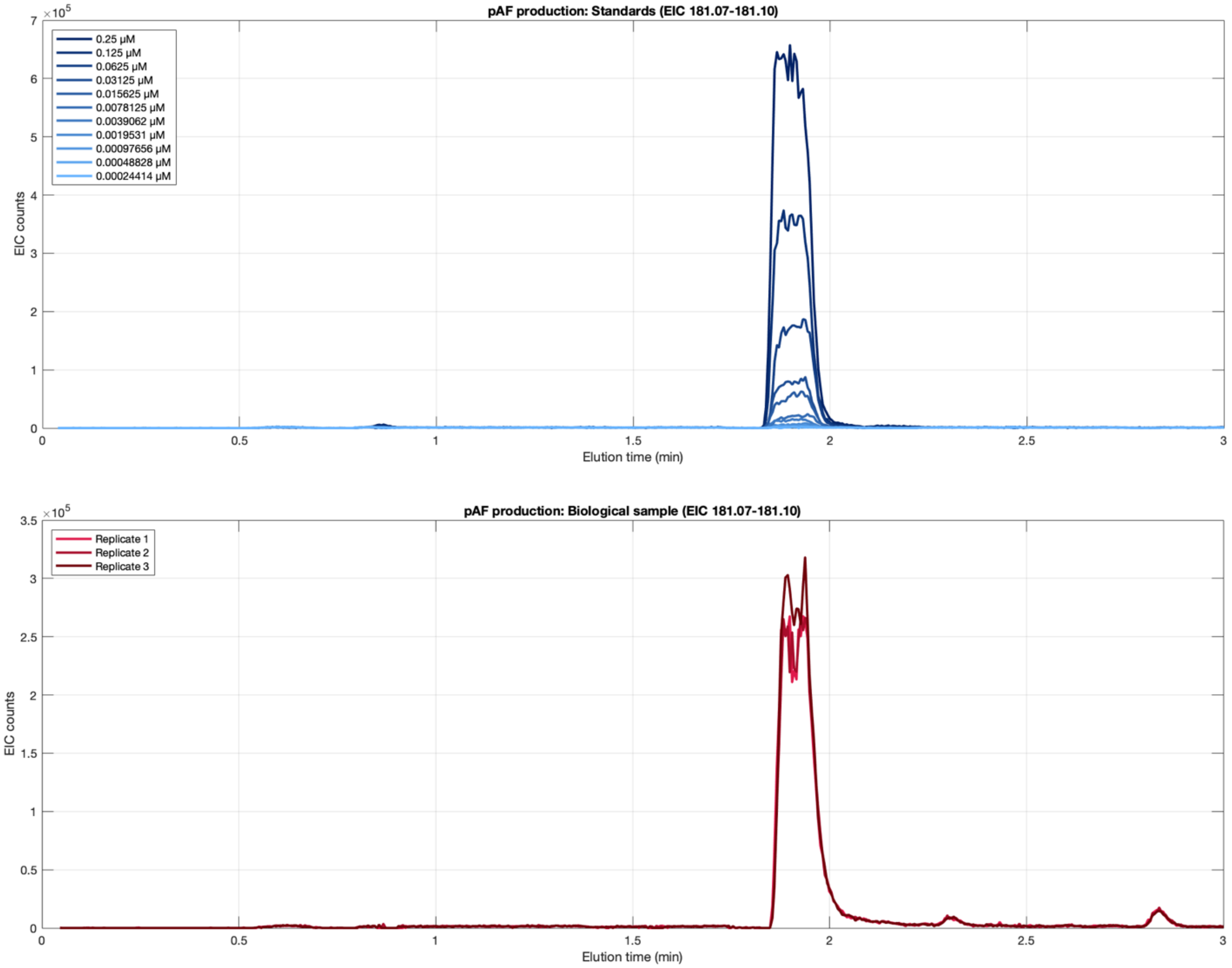


**Figure S4** – Detection of pAF by LC/MS. (top panel) Positive ion extracted ion chromatograms (EICs) (MW 181.07-181.10) for reference standards of pAF samples of known concentration. The reference standards were generated by adding pAF to the supernatant of Escherichia coli EcNR2 (non-pAF producing) cultures. (bottom panel) Representative EICs for supernatant collected from E. coli engineered to produce pAF through the heterologous expression of papBAC genes.

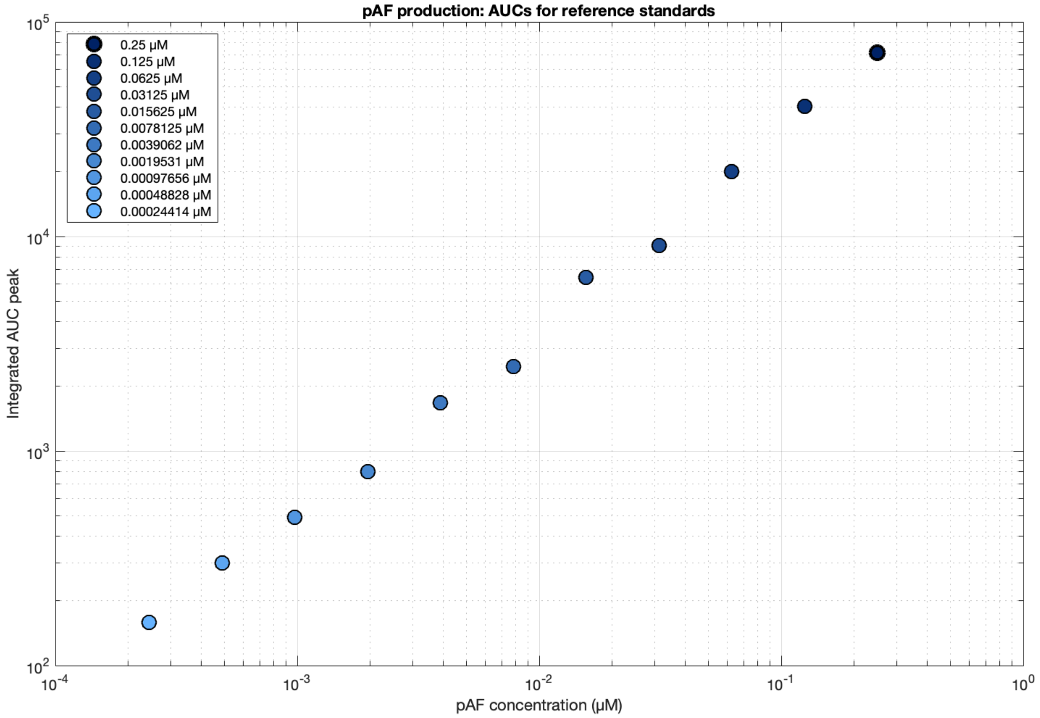


**Figure S5 –** Standard curve generated for pAF quantification by LC/MS. The area under the curve (AUC) of EIC peaks corresponding to pAF was calculated for reference standards of known concentration. A standard curve used to interpolate pAF concentrations in experimental samples was calculated through linear regression of the reference standard curve. Reference standards were generated in technical triplicate for each LC/MS experiment and the mean of the replicates was used for regression.
