## Supplementary Tables S1 and S2 for "Computational design and construction of an *Escherichia coli* strain engineered to produce a non-standard amino acid"

**Table S1.** Numerical data for the graph shown in main text Figure 3C.

| Strain | Plasmid | Measured doubling time (min) | Relative doubling time | [pAF] (μM) |
|---|---|---|---|---|
| ECNR2.ΔpheA | pPAF-v5 | 149 ± 1 | 1.084 ± 0.016 | 3.72 ± 0.32 |
| ECNR2.ΔtrpDE | pPAF-v5 | 143 ±0.02 | 1.048 ± 0.007 | 3.3 ± 1.1 |
| ECNR2.ΔtyrA | pPAF-v5 | 151 ± 0.4 | 1.099 ± 0.010 | 2.2 ± 1.0 |
| ECNR2.ΔpheA.ΔtrpDE | pPAF-v5 | 144 ± 5 | 1.052 ± 0.041 | 2.32 ± 0.17 |
| ECNR2.ΔtrpDE.ΔtyrA | pPAF-v5 | 165 ± 2 | 1.203 ± 0.023 | n.d. |
| ECNR2.ΔtyrA.ΔpheA | pPAF-v5 | 153 ± 2 | 1.116 ± 0.022 | n.d. |
| ECNR2.ΔpheA.ΔtrpDE.ΔtyrA | pPAF-v5 | 150 ± 1 | 1.093 ± 0.015 | 1.83 ± 0.19 |
| ECNR2.ΔpheA.ΔtrpDE.ΔtyrA.ΔentA | pPAF-v5 | 153 ± 0.6 | 1.113 ± 0.012 | n.d. |
| ECNR2 | pPAF-v5 | 137 ± 1 | 1.000 ± 0.014 | 3.95 ± 0.08 |
| ECNR2.aroFG-FBR | pPAF-v5 | 124 ± 5 | 0.907 ± 0.044 | 6.4 ± 2.2 |
| ECNR2.ΔtyrR | pPAF-v5 | 157 ± 3 | 1.145 ± 0.030 | n.d. |
| ECNR2.ΔtyrR.aroFG-FBR | pPAF-v5 | 161 ± 5 | 1.175 ± 0.043 | 36 ± 4.1 |
| ECNR2 | pGFP-RFP | 142 ± 2 | 1.038 ± 0.019 | n.d. |

**Table S2.** Vitamin and mineral supplements used in all culture conditions.

| **100x vitamin solution (for 500 ml in water)** | | |
|---|---|---|
| **Component** | **Concentration (M)** | **Volume (ml)** |
| Thiamine HCl | 0.02 | 25 |
| Calcium pantothenate | 0.02 | 25 |
| *p*-aminobenzoic acid | 0.02 | 25 |
| *p*-hydroxybenzoic acid | 0.02 | 25 |
| 2,3-dihydroxybenzoic acid | 0.02 | 25 |
| Water | | 375 |
| **50,000x trace metal mix (for 50 ml in water)** | | |
| **Component** | **Formula** | **Amount (g)** |
| Ammonium molybdate | $(NH_4)_6Mo_7O_{24} \bullet 4H_2O$ | 0.009 |
| Boric acid | $H_3BO_3$ | 0.062 |
| Cobalt chloride | $CoCl_2$ | 0.018 |
| Cupric sulfate | $CuSO_4$ | 0.006 |
| Manganese chloride | $MnCl_2$ | 0.04 |
| Zinc sulfate | $ZnSO_4$ | 0.007 |
