## Supplementary Software S2 for "Computational design and construction of an *Escherichia coli* strain engineered to produce a non-standard amino acid": suppl_software2.m

```
% Zomorrodi, Hemez, et al., Computational design and construction of an Escherichia coli strain engineered to produce a non-standard amino acid
% Supplementary Software S2 – MATLAB script used to calculate doubling times from plate reader growth curve data.

function [DT, R, median] = lnDoublingTime(ODvals, timevals, points, varargin)

% Check if the OD series is a filler dataset
if ODvals(1) == ODvals(end)
    DT = 0;
    R = 0;
    median = 0;
    return
end

% Check that the number of points for linear fitting is odd
if mod(points, 2) ~= 1
    error('Subset of points for linear regression must be odd')
end

% Check that the number of values in OD and time arrays is equal
if length(ODvals) ~= length(timevals)
    error('ODVALS and TIMEVALS must have the same number of values')
end

plotter = 0;
datapts = length(ODvals);

if nargin == 4
    if strcmp(varargin{1}, 'plot')
        disp('PLOT function active')
        plotter = 1;
        figure
    else
        error('Optional fourth argument is invalid')
    end
end

% Transform the data into logspace, transform inf to NaN
ODlog = log(ODvals);
ODlog(isinf(ODlog)) = NaN;
ODlog = real(ODlog);

% Perform linear regression to estimate doubling time
minRsq = 0.98; % R-squared value threshold
padding = floor(points/2);
maxSlp = 0; % Current highest slope
currVals = [0 0 0 0];

% currRsq: R-squared value of current highest slope

for i = (1+padding):(datapts-padding)
    Tmin = i - padding;
    Tmax = i + padding;
    currTvals = timevals(Tmin:Tmax);
    currODvals = ODlog(Tmin:Tmax);
    
    % Check if the region is above the linearity threshold
    currC = corrcoef(currTvals, currODvals);
    currR = currC(1,2);
    currRsq = currR^2;
    
    % If region is above linearity threshold, calculate the slope
    if currRsq >= minRsq
        linfit = polyfit(currTvals, currODvals, 1); % linfit: [slope, y-int]
        currSlp = linfit(1);
        
        % Update maximum slope; keep 
        if currSlp > maxSlp
            maxSlp = currSlp;
            Yint = linfit(2);
            currVals = [maxSlp Yint currRsq i];
        end
    end
    
end

maxVals = currVals;

% Do transformation on maximum values to get doubling time, centroid time
DT = log(2) / maxVals(1);
R = maxVals(3);
median = timevals(maxVals(4));

% Optional plotting function
if plotter == 1
    subplot(1,2,1)
    plot(timevals, ODvals, 'linewidth', 3)
    grid on
    xlabel('Time')
    ylabel('OD_{600}')
    title('Raw Data')
    
    subplot(1,2,2)
    hold on
    
    % Plot log-transformed data
    plot(timevals, ODlog, 'ok', 'markerfacecolor','k')
    
    % Plot the region of linear fit used to calculate doubling time
    fitTimes = timevals( (maxVals(4)-padding) : (maxVals(4)+padding) );
    fitLnOD = (fitTimes.*maxVals(1)) + maxVals(2);
    plot(fitTimes, fitLnOD, '-r', 'linewidth', 5)
    
    grid on
    xlabel('Time')
    ylabel('ln(OD_{600})')
    title('Log-Transformed Data')
end

end
```
